# Ultrastructural localization of calcium homeostasis modulator 1 in the taste buds of mouse circumvallate papillae

**DOI:** 10.1101/2023.09.15.557881

**Authors:** Rio Ikuta, Yuu Kakinohana, Shun Hamada

## Abstract

Taste receptor cells are morphologically classified as types II and III. These two types of cells transduce different taste modalities and form morphologically distinct synapses with afferent nerve fibers. Type III cells form conventional chemical synapses, whereas type II cells form a unique type of synapses referred to as channel synapses, which release adenosine triphosphate (ATP) as a neurotransmitter through voltage-gated channels comprising calcium homeostasis modulator (CALHM) 1 and CALHM3. Further, channel synapses are associated with large mitochondria with tubular cristae known as atypical mitochondria, which are believed to produce ATP for neurotransmission. Herein, we aimed to generate a monoclonal antibody against CALHM1 and examine its localization in taste buds using immunoelectron microscopy. CALHM1 was detected along the plasma membrane apposed to atypical mitochondria, indicating that CALHM1 is localized at the presynaptic membrane of channel synapses. Additionally, CALHM1 was detected along the plasma membrane lined by subsurface cisternae at sites apposed to afferent nerve fibers.

## Introduction

Taste buds are gustatory organs located predominantly in the epithelium of the lingual papillae and soft palate. Each taste bud contains 50–100 cells, which are morphologically classified into types I, II, III, and lV (Murray 1993; Murray et al. 1969). Among these, types II and III are taste receptor cells. Type II cells detect sweet, bitter, and umami tastes, whereas type III cells detect sour taste. Salty taste is likely to be received by both type II (Nomura et al. 2020; Ohmoto et al. 2020) and type III (Lewandowski et al. 2016; Oka et al. 2013; Roebber et al. 2019) cells. However, the exact cell types involved in salty taste sensation have not been fully elucidated.

Type II and III cells form synapses with afferent nerve fibers that transmit taste information (Chaudhari and Roper 2010; Roper and Chaudhari 2017). Type III cells form conventional chemical synapses with afferent nerve fibers, whereas type II cells form a unique type of synapses referred to as “channel synapses” that lack synaptic vesicles (Romanov et al. 2018; Taruno et al. 2021). At channel synapses, adenosine triphosphate (ATP) is released as a neurotransmitter from type II cells through a voltage-gated channel comprising a heterohexamer of calcium homeostasis modulator 1 (CALHM1) and CALHM3 (Ma et al. 2018; Taruno et al. 2021; Taruno et al. 2013).

Channel synapses are closely associated with mitochondria, which is another unique feature. The mitochondria in channel synapses are generally larger than typical mitochondria and have twisted tubular cristae. These mitochondria are closely apposed to the presynaptic membrane of channel synapses (Romanov et al. 2018). These characteristic mitochondria in type II cells have been observed using electron microscopy and are referred to as “atypical” mitochondria (Royer and Kinnamon 1988; Yang et al. 2012). Atypical mitochondria produce and retain large amounts of ATP for neurotransmission. Furthermore, they prevent a large amount of Ca^2+^ influx through CALHM1/3 channels, which are large-pore nonselective channels that allow Ca^2+^ influx into the cytosol (Romanov et al. 2018).

CALHM1 has been detected in channel synapses via immunohistochemistry (Kashio et al. 2019; Nomura et al. 2020; Romanov et al. 2018). Using super-resolution microscopy, CALHM1 has been detected at sites of contact between type II cells and afferent nerve fibers (Nomura et al. 2020; Romanov et al. 2018). The detected CALHM1-immnunoreactivity (IR) is accompanied by patches of cytochrome c, a mitochondrial marker (Romanov et al. 2018). Furthermore, immunoelectron microscopy using an enzyme-based detection system detected CALHM1-IR in and around the region of contact between the plasma membrane of type II cells and atypical mitochondria. Moreover, CALHM1-IR was not observed elsewhere in taste receptor cells (Romanov, et al., 2018). These observations indicate that CALHM1/3 channels are selectively localized at the presynaptic membrane of channel synapses.

Herein, we produced a monoclonal antibody against CALHM1 that could detect CALHM1 in sections treated with a mild epitope retrieval procedure, thereby allowing CALHM1 detection in structurally well-preserved tissues using immunoelectron microscopy. Using an immunogold-based detection system, we determined the localization of CALHM1 more accurately compared to the use of an enzyme-based detection system. Our results unequivocally showed that CALHM1 was localized in the presynaptic membrane of channel synapses. Further, CALHM1 was detected in the plasma membrane of type II cells lined by subsurface cisternae at the regions of contact between nerve fibers and type II cells, where CALHM1 has not been previously detected.

## Materials and methods

### Animals

For histological examination, 8-week-old male ICR mice were obtained from Japan SLC (Shizuoka, Japan). For monoclonal antibody production, three 8-week-old female Wistar rats (SLC) were used. The animals were housed with a standard laboratory diet and water under a 12-h light/dark cycle at 23°C ± 2°C. All experimental procedures were approved by the Animal Committee of Fukuoka Women’s University.

### Generation of CALHM1 monoclonal antibody

To obtain CALHM1 antibodies, we used the rat lymph node method (Kishiro et al. 1995). An antigenic peptide (Cys-PRKEVATYFSKV), corresponding to the 12 carboxyl- terminal residues of mouse CALHM1, was synthesized and conjugated with maleimide- activated hemocyanin (Imject™ Maleimide-Activated Blue Carrier Protein, Thermo Fisher Scientific, MA, USA). The conjugate solution was emulsified with Freund’s complete adjuvant (Becton, Dickinson and Company, NJ, USA) and injected into the base of the tail of Wistar female rats under isoflurane anesthesia. Two weeks after immunization, iliac lymph nodes were resected. Cells in the lymph nodes were collected, fused with P3U1 mouse myeloma cells using the usual method with polyethylene glycol, and seeded in 96-well plates in RPMI-1640 medium containing 10% fetal bovine serum, hypoxanthine, aminopterin, and thymidine. The wells containing hybridomas that produced anti-CALHM1 antibodies were identified using enzyme-linked immunosorbent assay for the antigenic peptide. Subsequently, hybridomas in the identified wells were cloned using the limiting dilution method. The supernatants of the identified hybridoma clones were used for immunofluorescence and immunoblotting to examine antibody specificity. Finally, the hybridoma clone 8F1 was selected.

### Cell culture, transfection, immunoblotting, and immunofluorescence of transfected cells

NIH3T3 cells were cultured in Dulbecco’s modified Eagle’s medium (DMEM) supplemented with 10% calf serum. Cells were incubated at 37°C in humidified 5% CO_2_/95% air. For immunofluorescence, transfected cells were cultured on poly L-lysine-coated coverslips.

To express CALHM1 in NIH3T3 cells, mouse CALHM1 cDNA in pcDNA 3.1(-), which was provided by Dr. Akiyuki Taruno, Kyoto Prefectural University of Medicine (Kashio et al. 2019), was transfected using HilyMax (Dojindo Laboratories, Kumamoto, Japan) according to the manufacturer’s instructions. The empty vector pcDNA3.1(−) was used for mock transfection. Transfected cells were cultured for 24–48 h and used for immunoblotting and immunofluorescence.

For immunoblotting, transfected cells were rinsed with phosphate-buffered saline (PBS) and lysed in 62.5 mM Tris-HCl (pH 6.8) containing 10% glycerol and 2% sodium dodecyl sulfate (SDS) with sonication. Next, 2-mercaptoethanol was added to the lysates at a concentration of 1%, and the lysates were boiled for 5 min. After adding bromophenol blue, the proteins were separated via 2% SDS-polyacrylamide gel electrophoresis and blotted onto nitrocellulose membranes. After blotting, the nitrocellulose membranes were rinsed briefly in PBS containing 0.3% TritonX-100 (PBST) and incubated in 5% skim milk diluted with PBST for 1 h. The membrane was washed with PBST and incubated in 8F1 diluted in PBST containing 5% blocking reagent (Blocking One; Nacalai Tesque, Kyoto, Japan) for 1 h at room temperature. After washing with PBST, the membranes were incubated with peroxidase-conjugated goat anti-rat antibody (1:50000, Jackson ImmunoResearch, PA, USA) diluted with PBST containing 5% Blocking One for 1 h at room temperature. After washing with PBST, the binding sites of the secondary antibodies were detected using Western Lightning ECL Pro (PerkinElmer, MA, USA). Chemiluminescence was visualized using the Fusion Solo S imaging system (Vilber Bio Imaging, Paris, France).

For immunofluorescence, transfected cells on coverslips were rinsed with DMEM and fixed with methanol for 20 min on ice. After fixation, methanol was aspirated, and the coverslips were dried. After rinsing with PBS, the cells were incubated in Blocking One solution containing 5% normal goat serum for 30 min at room temperature. After rinsing with PBS, coverslips were incubated with a mixed solution of 8F1 and 32C2, a commercially available monoclonal antibody against CALHM1, overnight at 4°C. Table 1 shows the primary and secondary antibodies used for immunofluorescence and immunoelectron microscopy. After rinsing twice with PBST for 5 min each, the cells were incubated with Alexa 488-labeled goat anti-rat antibody diluted in PBS containing 5% Blocking One for 1 h at room temperature. After rinsing with PBST, the cells were incubated with Alexa-594- labeled goat anti-mouse antibody for 1 h. After rinsing with PBST and PBS, the nuclei were labeled with 4’,6-diamidino-2-phenylindole (DAPI, 1 μg/mL) and then coverslipped with a mounting medium (Fluoroshield Mounting Medium; ImmunoBioScience, CA, USA). Images were obtained using a fluorescence microscope (Eclipse E800; Nikon, Tokyo, Japan) equipped with the DS-Fi3 digital camera system (Nikon).

**Table 1.**
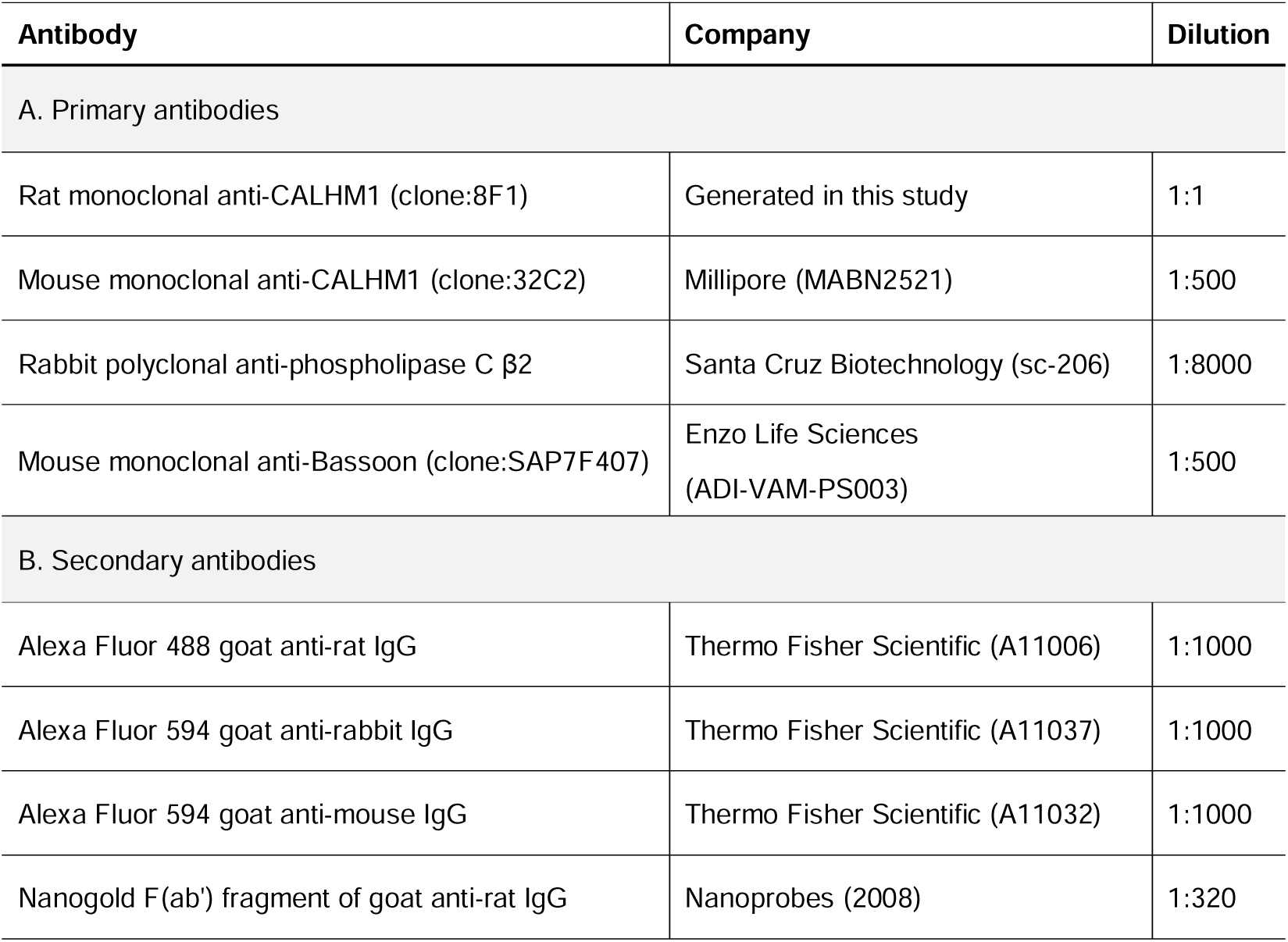
Antibodies used in the present study.

### Immunofluorescence of the lingual papillae

Mice were deeply anesthetized with sodium pentobarbital (Somnopentyl; Kyoritsu Seiyaku, Tokyo, Japan) and perfused through the ascending aorta with saline, followed by fixatives for 10 min. The fixative was 2% paraformaldehyde in 0.1 M phosphate buffer (PB, pH 7.3) for light microscopy and periodate-lysine-paraformaldehyde fixative (McLean and Nakane 1974) for immunoelectron microscopy. After perfusion fixation, tissues including the circumvallate or fungiform papillae were removed under a binocular microscope and post- fixed in the same fixative at 4°C for 2 h. The tissues were then immersed in 25% sucrose in 0.1 M PB at 4°C overnight for cryoprotection and then frozen in embedding medium (Tissue-Tek OCT Compound; Sakura FineTek Japan, Tokyo) with hexane cooled with dry ice. The frozen tissues were stored at −80°C for later analyses.

For immunofluorescence, 7-μm-thick sections were cut using a cryostat, collected on silane-coated slides, and air-dried for 30 min. The sections were treated with 10 mM citrate buffer (pH 7.0) for 20 min at 85°C for antigen retrieval. The sections were then rinsed in PBST and treated with Blocking One solution containing 3% MOM Blocking Reagent (Vector, CA, USA) for 1 h. After rinsing briefly with PBS, the sections were incubated with a mixture of primary antibodies (Table 1) diluted in PBS containing 5% Blocking One at 4°C overnight. After incubation with primary antibodies, the sections were rinsed with PBST and PBS. The sections were then incubated with the appropriate secondary antibodies (Table 1) labeled with Alexa 488 at room temperature for 1 h. After rinsing with PBST and PBS, secondary antibodies labeled with Alexa 594 were applied to the sections for 1 h. The sections were then rinsed with PBST and PBS, incubated with DAPI solution, and coverslipped with a mounting medium. Controls were prepared by omitting the primary antibodies. Images were obtained using a confocal laser microscope (C2 Plus; Nikon). The contrast and brightness of the images were adjusted using Adobe Photoshop software.

### Immunoelectron microscopy

Frozen tissues fixed with periodate-lysine-paraformaldehyde fixative were cut into 10- µm-thick sections, collected on silane-coated slides, and air-dried for 15 min. The sections were incubated in 10 mM citrate buffer (pH 7.0) for 20 min at 40°C for epitope retrieval. After rinsing with PBS for 5 min, the sections were blocked with PBS containing 20% Blocking One and 0.005% saponin for 15 min. The sections were incubated with the 8F1 hybridoma culture supernatant at 4°C for 42–48 h. After rinsing with PBS containing 0.005% saponin, the sections were incubated with the F(ab’) fragment of goat anti-rat IgG labeled with 1.4 nm colloidal gold (Table 1) at 4°C overnight. After rinsing four times with PBS containing 0.005% saponin for 10 min each, the sections were rinsed with 0.1 M PB to remove saponin. The sections were then fixed with 1% glutaraldehyde in 0.1 M PB for 10 min. After rinsing with 0.1 M PB, the sections were rinsed with 50 mM HEPES (pH 5.8) three times for 5 min each.

The binding sites of the gold-labeled antibodies were silver-enhanced using a kit (HQ Silver; Nanoprobes, NY, USA) at 20°C in the dark and washed twice with water. The sections were post-fixed with 0.5% osmium tetroxide in 0.1 M PB at 4°C for 1.5 h. After rinsing with water, the sections were dehydrated by passing through a graded series of ethanol (50%, 70%, 80%, 95%, 99.5%, and 100%) and propylene oxide and then embedded in epoxy resin (TAAB Laboratories, UK). Ultrathin sections (80–90 nm) were cut using an ultramicrotome (EM UC7; Leica, Germany), collected on grids, stained with uranyl acetate and lead citrate, and observed under an electron microscope (JEM-1400 Plus; JEOL, Tokyo, Japan). The contrast of the images was adjusted using Adobe Photoshop software.

## Results

To investigate the localization of CALHM1, we first used 32C2, a commercially available mouse monoclonal antibody against CALHM1, which has been previously used for the immunohistochemical analysis of taste buds (Romanov et al. 2018). However, this antibody barely detected CALHM1 in tissues treated with our immunoelectron microscopy experimental procedures. Therefore, we produced 8F1, a rat monoclonal antibody, which specifically recognizes the C-terminal region of mouse CALHM1.

Immunoblotting using 8F1 against NIH3T3 cell lysates transfected with an expression vector encoding mouse CALHM1 cDNA yielded a single band of approximately 40 kDa, which was consistent with the calculated molecular weight of mouse CALHM1 (38.26 kDa) (Figure 1 a). No bands were detected in the lysates of the mock-transfected cells.

**Figure 1.**
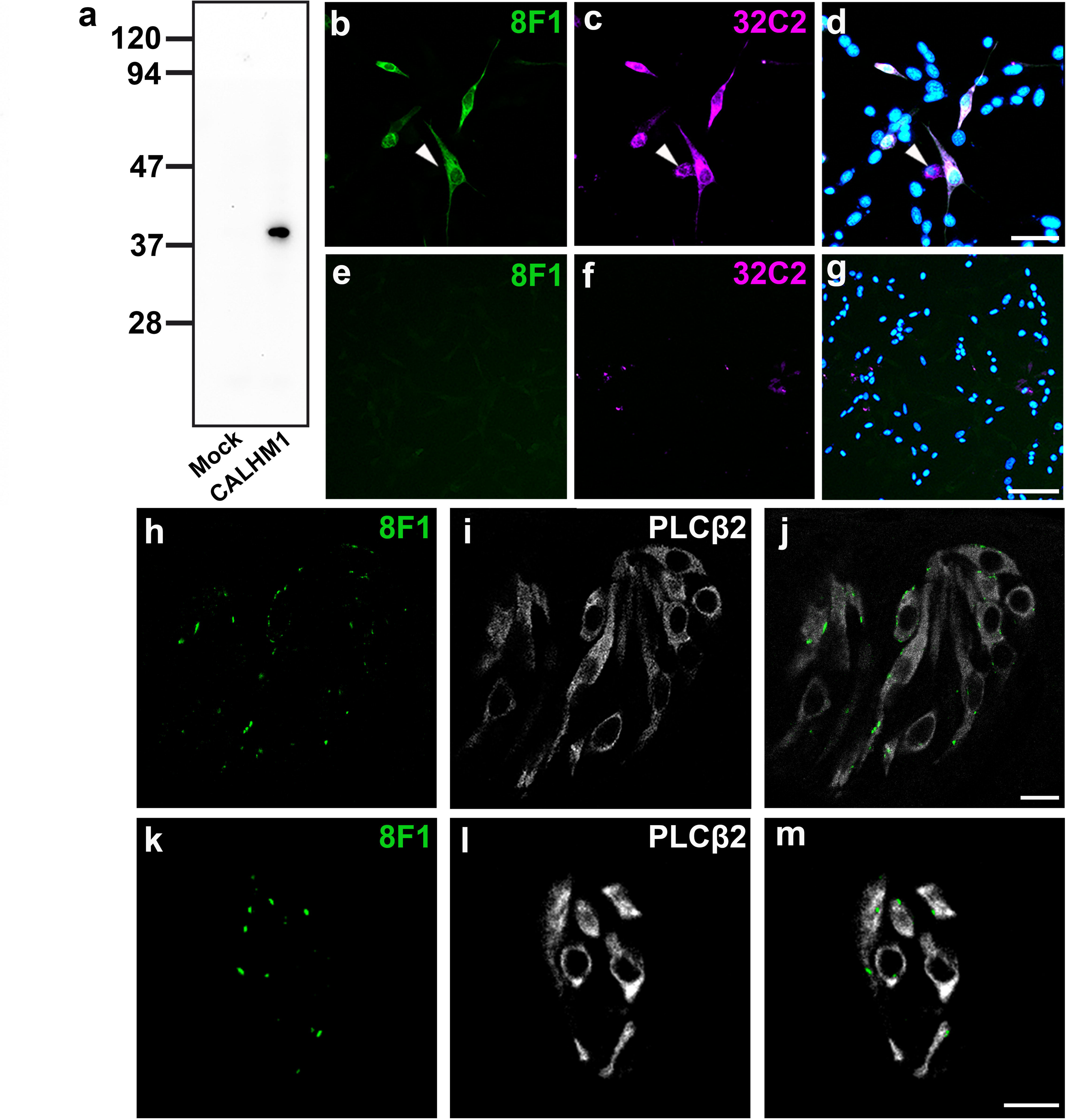
Characterization of rat monoclonal antibody 8F1 against CALHM1. Immunoblotting of whole-cell lysates obtained from NIH3T3 cells transfected with mouse CALHM1 cDNA or empty vector using the rat monoclonal antibody 8F1 (**a**). NIH3T3 cells expressing mouse CALHM1 were double-immunolabeled with 8F1 (**b**, green) and commercially available CALHM1 antibody 32C2 (**c**, magenta). Merged images of **b** and **c** with DAPI-stained nuclei (**d**). The 8F1 and 32C2 antibodies labeled almost the same cells. However, a small subset of transfected cells was labeled only with 32C2 (**b**–**d**, arrowheads). Mock-transfected NIH3T3 cells were double-immunolabeled with 8F1 (**e**, green) and 32C2 (**f**, magenta) antibodies. Merged images of **e** and **f** with DAPI-stained nuclei (**g**). A small subset of mock-transfected NIH3T3 cells was labeled with 32C2 but not with 8F1 antibody (**e**–**g**). Representative confocal images (**h**–**j**, individual Z-sections) of taste buds in circumvallate papillae double-immunolabeled with 8F1 (**h**) and PLCβ2 (**i**) antibodies. Merged images of **h** and **i** (**j**). 8F1-IR was observed in the peripheral regions of PLCβ2-positive cells. Representative confocal images (**k**–**m**, individual Z-sections) of taste buds in fungiform papillae double-immunolabeled with 8F1 (**k**) and PLCβ2 (**l**) antibodies. Merged images of **k** and **l** (**m**). The distribution pattern of 8F1-IR in the taste buds of fungiform papillae was similar to that of circumvallate papillae. Scale bars = 50 μm (**d**), 100 μm (**g**), and 10 μm (**j**, **m**).

Next, we performed double immunolabeling with 8F1 and 32C2 in CALHM1- transfected NIH3T3 cells. These two monoclonal antibodies labeled nearly identical cells; however, some cells were labeled with 32C2, but not with 8F1 (Figure 1 b–d). Furthermore, 32C2 signals were observed in mock-transfected NIH3T3 cells (Figure 1 f), thereby indicating that 32C2 may react with endogenous molecules in NIH3T3 cells, other than CALHM1.

We used 8F1 to detect endogenous CALHM1 in the mouse taste buds. 8F1-IR was observed as small puncta in the taste buds of circumvallate papillae. 8F1 immunoreactive puncta were observed along the cellular profiles expressing phospholipase C β2 (PLCβ2), a marker of type II cells (Figure 1 h–j). The same distribution pattern was observed in the taste buds of fungiform papillae (Figure 1 k–m). These distribution patterns of 8F1-IR were consistent with those of CALHM1-IR with other antibodies (Kashio et al. 2019; Nomura et al. 2020; Romanov et al. 2018).

### Localization of CALHM1 at channel synapses

8F1 detected CALHM1 in taste bud tissues treated with a mild epitope retrieval procedure. Thus, 8F1 was used to detect CALHM1 via immunoelectron microscopy using the pre-embedding immunogold-silver labeling technique. As expected, silver deposits showing CALHM1-IR were mainly observed in the restricted regions along the plasma membrane of the cells with ultrastructural features of type II cells: an oval nucleus with an electron-translucent cytoplasm (Figure 2 a). In addition to the plasma membrane, silver deposits were occasionally observed in multivesicular bodies (data not shown).

**Figure 2.**
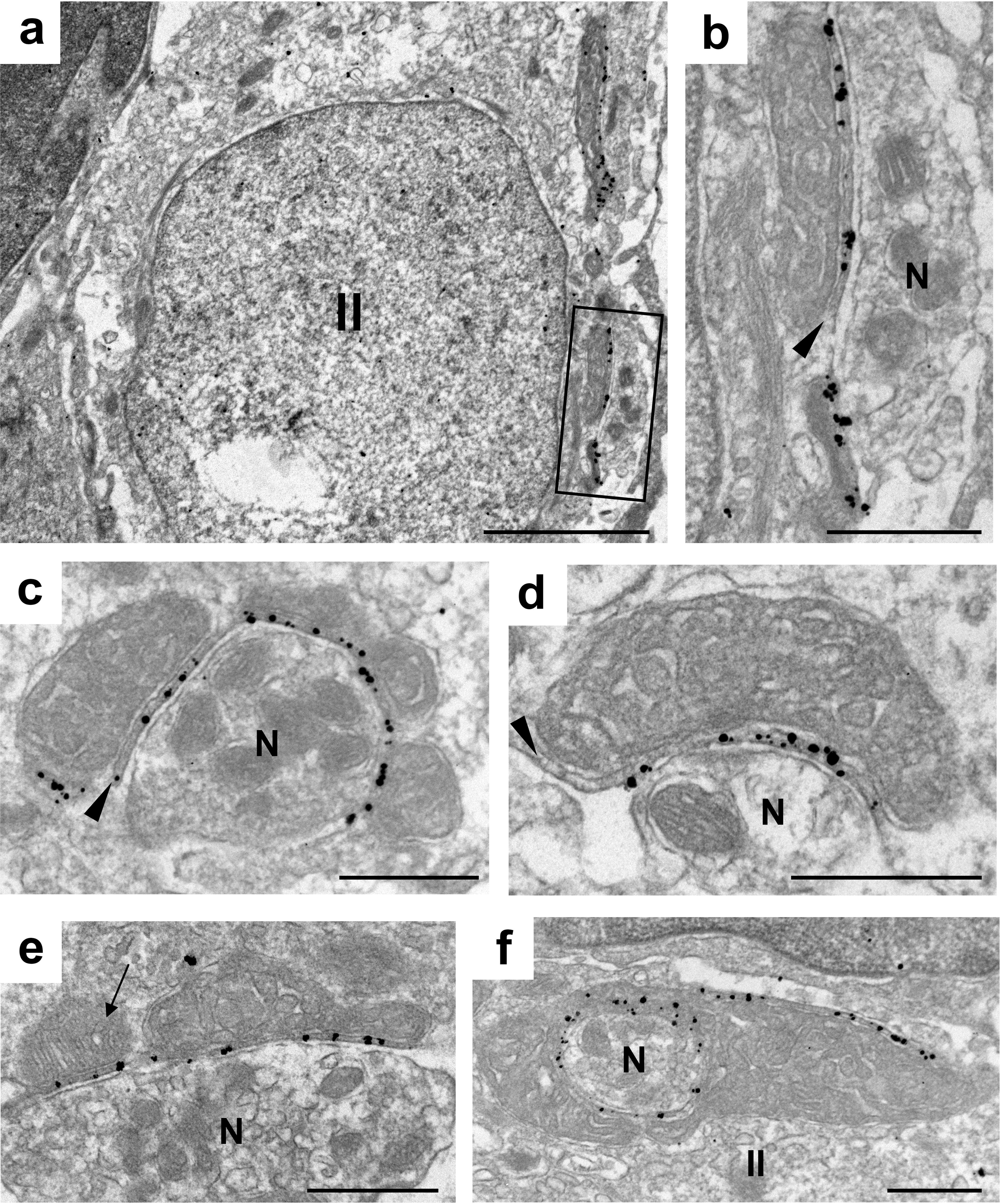
Representative immunoelectron micrographs showing the localization of CALHM1-IR in the plasma membrane apposed to mitochondria. Silver deposits of CALHM1-IR were detected along the plasma membrane apposed to atypical mitochondria at the sites of contact between type II cells and nerve fibers (**a**). A higher-magnification image of the rectangular area shown in **a** (**b**). Electron-dense material (arrowheads) was observed between the outer membrane of atypical mitochondria and the plasma membrane with CALHM1-IR (**b**-**d**). Atypical mitochondria surrounding the entire circumference of nerve fibers that invaginate into type II cells were occasionally observed (**e**). CALHM1-IR was also detected along the plasma membrane apposed to the mitochondria with lamellar cristae (**f**, arrow). Type II cells and afferent nerve fibers are labeled as “II” and “N,” respectively. Scale bars = 2 μm (**a**) and 500 nm (**b**– **f**).

Membrane regions with silver deposits were often in close apposition to mitochondria at sites of contact with nerve fibers (Figure 2 b). Most, but not all, of these mitochondria are considered atypical. That is, they are large and have enlarged tubular cristae, indicating that CALHM1-IR was observed along the presynaptic membrane of channel synapses. As previously demonstrated using conventional electron microscopy (Royer and Kinnamon 1988), electron-dense material was observed between the presynaptic membrane and outer membrane of atypical mitochondria (Figure 2 b-d, arrowheads). Silver deposits of CALHM1- IR in channel synapses were predominantly detected along the plasma membrane lined by electron-dense material. Occasionally, we encountered channel synapses with atypical mitochondria surrounding the entire circumference of nerve fibers that invaginated into type II cells (Figure 2 e). These channel synapses are reminiscent of “fingerlike synapses,” a subset of synapses previously found in type III cells (Kinnamon et al. 1985; Royer and Kinnamon 1988).

In addition to atypical mitochondria, mitochondria with lamellar cristae were closely apposed to the plasma membrane at sites of contact with nerve fibers. Some mitochondria were in proximity to the atypical mitochondria (Figure 2 f). CALHM1 was detected along the plasma membrane apposed to these mitochondria with lamellar cristae as well as atypical mitochondria (Figure 2 f).

### CALHM1 is localized at the plasma membrane lined by subsurface cisternae

The subsurface cisterna is a flattened endoplasmic reticulum closely apposed to the plasma membrane and is mainly observed in neurons, sensory cells, and muscle cells (Chen et al. 2019; Rosenbluth 1962). In type II cells, subsurface cisternae have been observed in proximity to the plasma membrane facing nerve fibers. However, their role in taste transduction remains elusive (Clapp et al. 2004; Fujimoto and Murray 1970; Hisatsune et al. 2007; Royer and Kinnamon 1988; Takeda 1976; Yoshie et al. 1990).

Surprisingly, we detected CALHM1-IR in the plasma membrane that was closely apposed to subsurface cisternae in type II cells, in addition to channel synapses (Figure 3). As observed between atypical mitochondria and the presynaptic membrane of channel synapses, the narrow gap between the subsurface cisternae and the plasma membrane was filled with electron-dense material (Figure 3 b–d, arrowheads). Furthermore, we encountered subsurface cisternae surrounding the circumference of nerve fibers that invaginated into type II cells (Figure 3 e), as is the case with channel synapses (Figure 2 e). The subsurface cisternae were occasionally accompanied by atypical mitochondria (Figure 3f, g), as reported previously (Royer and Kinnamon 1988).

**Figure 3.**
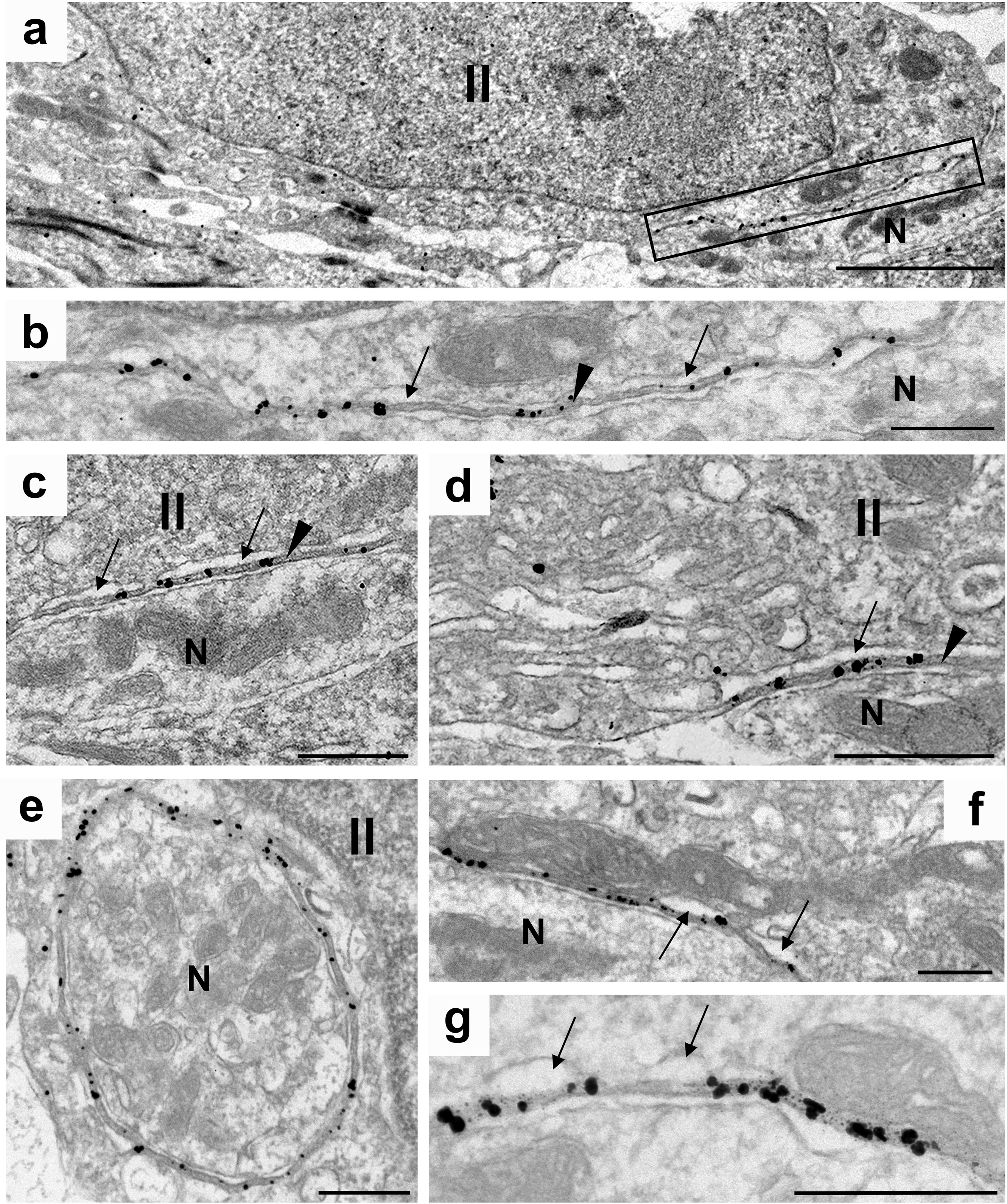
Representative immunoelectron micrographs showing the localization of CALHM1-IR in the plasma membrane apposed to subsurface cisternae. CALHM1 was detected in the plasma membrane that was not apposed to atypical mitochondria (**a**). A higher- magnification image of the rectangular area shown in **a** (**b**). CALHM1 was detected along the plasma membrane apposed to the subsurface cisternae (**b**–**d**, arrows). The space between the subsurface cisternae and plasma membrane was filled with electron-dense material (**b**–**d**, arrowheads). As in the case with the atypical mitochondria shown in Figure 2f, the subsurface cisternae occasionally surround the entire circumference of nerve fibers that invaginate into type II cells (**e**). We also encountered a few regions where CALHM1 was localized along the plasma membrane apposed to both the atypical mitochondria and subsurface cisternae (**f** and **g**). Type II cells and afferent nerve fibers are labeled as “II” and “N,” respectively. Scale bars = 2 μm (**a**) and 500 nm (**b**–**g**).

### CALHM1 was not detected in the synapses of type III cells

Synaptic transmission between type III cells and nerve fibers is mediated by conventional chemical synapses. However, a subset of these type III cell synapses is accompanied by mitochondria apposed to the plasma membrane, as is the case with atypical mitochondria in the channel synapses of type II cells. Thus, this subset of type III cell synapses has been referred to as “mixed” or “hybrid” synapses in previous studies (Wilson et al. 2022; Yang et al. 2020). However, whether the mitochondria in these hybrid synapses function as atypical mitochondria in the channel synapses of type II cells remains unclear.

Using immunoelectron microscopy, we could not detect CALHM1-IR in hybrid synapses (six hybrid synapses in two mice), even in a hybrid synapse located near a CALHM1-positive channel synapse (Figure 4 a–c). To further examine the absence of CALHM1 in hybrid synapses, we performed double immunofluorescence with CALHM1 and Bassoon, a marker of conventional synapses in type III cells (Ikuta and Hamada 2022) (Figure 4 d–f). Colocalization of CALHM1 and Bassoon was not detected in taste buds (67 taste bud profiles in three mice). These results suggest that CALHM1 is not localized at hybrid synapses.

**Figure 4.**
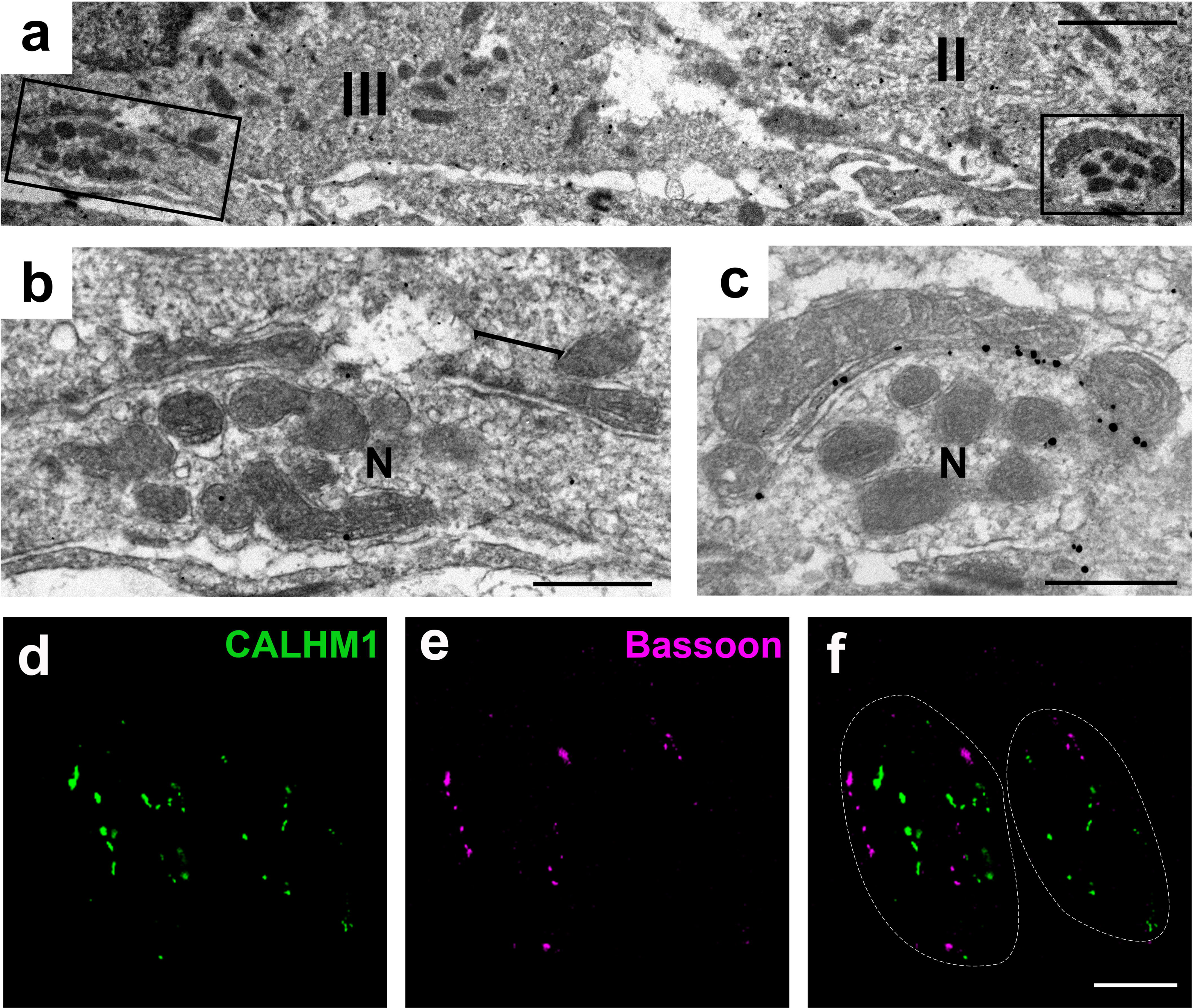
CALHM1 was not detected in the hybrid synapses of type III cells. **b** and **c** are high- magnification images of the left and right rectangular areas in **a**, respectively. CALHM1 was rarely detected along the plasma membrane apposed to the atypical mitochondria of hybrid synapses in type III cells (**b**). CALHM1 was detected in a channel synapse near a hybrid synapse shown in **b** (**c**). The bar in **b** indicates the active zone of the hybrid synapse. Double immunolabeling of CALHM1 (**d**, green) and Bassoon (**e**, magenta), a type III cell synapse marker. Merged images of **d** and **e** (**f**). Colocalization of CALHM1 and Bassoon was not detected. Type II cells, type III cells, and nerve fibers are labeled as “II,” “III,” and “N,” respectively. Scale bars = 2 μm (**a**), 500 nm (**b**, **c**), and 10 μm (**f**).

## Discussion

A previous study has examined the ultrastructural localization of CALHM1 via immunoelectron microscopy using an enzyme-based detection system combined with heat- induced epitope retrieval (Romanov et al. 2018). In general, the disadvantage of enzyme- based detection systems in immunoelectron microscopy is the diffusion of signals that obscures the exact localization site of the antigen. Moreover, the heat-induced epitope retrieval procedure used in a previous study (Romanov et al. 2018) may impair the ultrastructure of the tissues. We developed a monoclonal antibody that can detect the location of CALHM1 using an immunogold-based detection method in combination with moderate epitope retrieval, which facilitated a more accurate examination of the distribution of CALHM1 in well-preserved tissue structures. We unequivocally showed that silver deposits displaying CALHM1-IR were gathered along the plasma membrane closely apposed to the outer membrane of atypical mitochondria at sites of contact between nerve fibers and type II cells. Our results support the validity of a previously proposed structural model for channel synapses (Romanov et al. 2018).

As previously demonstrated using conventional electron microscopy (Royer and Kinnamon 1988), the space between the plasma membrane and atypical mitochondria is filled with electron-dense material. This finding indicates that the space between the plasma membrane and atypical mitochondria is filled with high-density protein molecules, which may form conduits of ATP and barriers for Ca^2+^ entry into the cytoplasm through CALHM1/3 channels, with scaffolding proteins.

CALHM1 was also detected along the plasma membrane apposed to mitochondria with lamellar cristate at regions of contact between type II cells and nerve fibers (Figure 2 f). These mitochondria with lamellar cristae may be mitochondria in transition between conventional and atypical, as previously described using serial block-face scanning electron microscopy (Wilson et al. 2022). Alternatively, the morphology of cristae in the mitochondria of channel synapses may change rapidly in response to synaptic activity or other factors. Nevertheless, our results showed that channel synapses are not always equipped with atypical mitochondria with tubular cristae.

In addition to channel synapses, we unexpectedly detected CALHM1 at the plasma membrane apposed to subsurface cisternae at sites of contact with nerve fibers. Subsurface cisternae are predominantly found at close appositions between type II cells and nerve fibers (Clapp et al. 2004; Royer and Kinnamon 1988; Takeda 1976; Yoshie et al. 1990). However, their exact function remains unknown. The present study revealed that the characteristics of the subsurface cisternae and plasma membrane apposed to them in type II cells were similar to those of channel synapses, including the localization of CALHM1, electron-dense material in the gap between the plasma membrane and subsurface cisternae, and a topographical relationship with nerve fibers (Figures 2 e and 3 e). Based on analogy with channel synapses, we speculated that subsurface cisternae might store and release ATP through CALHM1/3 channels. Because type II cells express the vesicular nucleotide transporter (VNUT) (Iwatsuki et al. 2009), VNUT may transport ATP into subsurface cisternae. Another possibility is that subsurface cisternae are transiently formed during the formation or degeneration of channel synapses to prevent the leakage of small molecules, including ATP, and the diffusion of Ca^2+^ into the cytoplasm through the large pores of CALHM1/3 channels. Observations by us and others (Royer and Kinnamon 1988) that subsurface cisternae are occasionally accompanied by atypical mitochondria may support this possibility.

Mitochondria similar to atypical mitochondria in type II cells have been observed in a subset of type III cells. These mitochondria have tubular cristae and are apposed to the plasma membrane at the sites of contact with nerve fibers. Nevertheless, they are more likely to be smaller than atypical mitochondria in type II cells. These atypical-like mitochondria in type III cells are found in proximity to conventional chemical synapses. Thus, they may be involved in the synaptic transmission of type III cells and are referred to as mixed or hybrid synapses (Wilson et al. 2022; Yang et al. 2020). However, we could not detect CALHM1 along the plasma membrane apposed to atypical-like mitochondria in type III cells using immunoelectron microscopy. Furthermore, we could not detect the colocalization of CALHM1 and Bassoon, a marker of the synapses of type III cells (Ikuta and Hamada 2022). Our results are consistent with those of a previous study showing that the expression of CALHM1 is confined to type II cells (Taruno et al. 2013). Thus, atypical-like mitochondria in type III cells do not form channel synapses and may be indirectly involved in the synaptic transmission of type III cells.

## Author Contribution

Study concept and design: S.H.; acquisition of data: R.I., Y.K.; analysis and interpretation of data; R.I., S.H.; drafting of the manuscript: R.I., S.H.; obtained funding: R.I., S.H.

## Ethical approval

The care and use of animals were approved by the Institutional Animal Care and Use Committee of Fukuoka Women’s University, in compliance with the Guidelines for Proper Conduct of Animal Experiments (Science Council of Japan).

## Data availability

The data that support the findings of this study are available from the corresponding author upon reasonable request.

